# Yield, nutritional quality, and fatty acid content of organic winter rye (*Secale cereale*) and winter wheat (*Triticum aestivum*) forages under cattle (*Bos taurus*) grazing conditions

**DOI:** 10.1101/688952

**Authors:** Hannah N. Phillips, Bradley J. Heins, Kathleen Delate, Robert Turnbull

**Affiliations:** West Central Research and Outreach Center, University of Minnesota, Morris, Minnesota, United States of America; Department of Animal Science, University of Minnesota, St. Paul, Minnesota, United States of America; Departments of Agronomy and Horticulture, Iowa State University, Ames, Iowa, United States of America

## Abstract

The objective of this study was to assess yield, nutritional quality, and fatty acid compositions of winter rye (*Secale cereale*) and winter wheat (*Triticum aestivum*) forages for grazing cattle (*Bos taurus*) in an organic system. The rye and wheat were established on two 4 ha plots in September 2015. Six groups of dairy steers rotationally grazed rye (n = 3) or wheat (n = 3) plots divided into seven paddocks (n = 14) from April to June 2016. Forage samples (n = 96) taken prior to paddock grazing were used to analyze forage characteristics. Mixed models with fixed factors of forage, date, and their interaction, a random subject factor of group nested in paddock, and a repeated effect of date were used for each outcome. The linear effect of date on fatty acids was obtained by substituting date as a continuous variable. The mean forage yield for rye was greater (*P* < 0.05) than wheat (mean ± standard error; 2840 and 2571 ± 82 kg ha^-1^, respectively). However, rye yielded less in the latter part of the grazing period. Wheat (19.3 ± 0.30% DM) had greater (*P* < 0.001) crude protein than rye (17.6 ± 0.30% DM). In general, crude protein, digestibility, and minerals decreased during the grazing period. Wheat (66.3 ± 0.54 g 100g^-1^) had greater (*P* < 0.001) alpha-linolenic acid (18:3*n*-3) concentration than rye (63.3 ± 0.54 g 100g^-1^). Although both forages decreased (*P* < 0.05) in alpha-linolenic acid concentration, wheat decreased 2.49 times more (*P* < 0.001) per d compared to rye forage. Winter rye and winter wheat forages are viable for cattle grazing. Producers should initiate early grazing to maximize protein, digestibility, and alpha-linolenic acid intake while the forages are immature.

## Introduction

The growing demand for organic beef and dairy products [1] is partially driven by consumer interest in perceived health benefits [2] of the altered fatty acid (**FA**) profiles of organic beef and milk lipids [3–6]. Products from organic cattle have a desirable FA profile, including greater concentrations of conjugated linoleic acid and omega-3 FA (***n*-3**), and lower omega-6:omega-3 FA ratios (***n*-6:*n*-3**), compared to conventionally-fed cattle that consume a greater proportion of their diet as grain and grain-derived feedstuff [3,4,6]. A greater concentration of alpha-linolenic acid (**18:3*n*-3**) in forages has a positive influence on the concentration of *n*-3 in milk and lipids of beef [7]. Furthermore, fresh forages contain even greater concentrations of 18:3*n*-3 than processed or stored forages [8, 9]. Thus, increasing the intake of fresh forages via pasture grazing can improve the nutritional quality of milk and beef products while allowing producers an opportunity to capitalize on forage production for grazing systems.

The rules of the United States Department of Agriculture National Organic Program (§205.237) [10] require that cattle consume at least 30% of their dry matter intake from pasture, except during the finishing phase for beef, and require an active soil building plan to limit soil erosion and nutrient leaching. Pasture grazing is also a low-cost method to feed cattle compared to feeding stored organic grains and forages [11]. Hence, one of the main obstacles organic beef and dairy producers face is lack of pasture forages for grazing [12]. In the upper Midwest of the United States of America (**USA**), perennial grasses and legumes, such as alfalfa (*Medicago sativa*), chicory (*Cichorium intybus*), meadow brome grass (*Bromus biebersteinii*), meadow fescue (*Festuca pratensis*), orchard grass (*Dactylis glomerata*), perennial ryegrass (*Lolium perenne*), red clover (*Trifolium pratense*), and white clover (*Trifolium repens*), are common pasture forages typically grazed beginning in May. Establishing cold hardy winter cover crops in the fall to protect bare soil after crop harvest is a popular method to meet the soil building requirement [13]. Furthermore, these winter cover crops yield forage in early spring; thus, winter cover crop grazing — prior to perennial pasture grazing in the spring — may be advantageous for producers by extending the grazing season [14]. Winter rye (*Secale cereale*; **WR**) and winter wheat (*Triticum aestivum*; **WW**) are adapted to low temperatures and yield forage in the early spring. Although, integrating small grain cover crops into pasture systems extends the grazing season, farmers may be reluctant to graze livestock on the forages because of inconstant nutritional quality and rapid decreases in nutritional quality as forages mature [15].

As pasture-based beef and dairy industries grow, it is important to assess alternative forages and understand their impacts on production and nutritional quality for grazing. Therefore, the objectives of this study were to assess and compare WR and WW pastures for forage yield, dry matter (**DM**), nutritional quality, mineral composition, and FAs during the grazing season.

## Materials and methods

### Ethical statement

Researchers conducted this study at the University of Minnesota West Central Research and Outreach Center (**WCROC**) in Morris, Minnesota, USA. The University of Minnesota Institutional Animal Care and Use Committee approved all animal care and management (Protocol Number: 1411-32060A).

### Study background

The WCROC research dairy has 300 low-input conventional and organic grazing cows. The organic herd was certified with the Midwest Organic Services Association in June 2010, which is accredited by the United States Department of Agriculture National Organic Program. The herd was part of a crossbreeding program established in 2000, as described by Heins et al. [16]. A sociological component of the current study detailed specific obstacles related to integrating crops and livestock as identified by producers over the course of the project, and reported increasing support for integrated crop-livestock systems resulting in growing communities of practice in which farmer-to-farmer knowledge exchange and peer support overcome obstacles to success in these systems [17].

### Pasture establishment

The WR (*S. cereale*) and WW (*T. aestivum*) were established on two adjacent 4 ha plots in September 2015. The study chose these forages based on their hardiness and popularity as cover crops in the upper Midwest of the USA. Prior to planting, the WCROC utilized the plots for grazing dairy cattle and included perennial forages for at least 20 years. Manure from cattle deposited during grazing fertilized pastures, and the study used no additional fertilizer or irrigation.

### Experimental approach

Six groups of five Holstein and crossbred dairy steers (*Bos taurus*; n = 30) born at the WCROC (March – May 2015) were used in this study. Steers were grouped by age and breed composition. Details on care are explained by Phillips et al. [18]. Prior to grazing, steers were housed in a loose-confinement barn in their respective groups, and were fed an organic total mixed ration diet of corn silage, alfalfa silage, corn, soybean meal, and minerals from weaning (ca. 10 weeks of age) until grazing (ca. 12 months of age). One steer died from peritonitis and was removed from the study prior to grazing.

Grazing initiated when forage height reached 15 cm in plots on 25 April 2016. The six steer groups were randomly assigned to graze WR (n = 3) or WW (n = 3), and forages were balanced by age and breed. Steers remained in their groups throughout the grazing period and were separated by temporary electric fencing. The groups rotationally grazed 0.57 ha paddocks in WR (n = 7) and WW (n = 7) plots for seven weeks, supplemented by ad libitum minerals. The stocking density was approximately 25 ha^-1^ (10,650 kg ha^-1^) per paddock. Groups rotated paddocks every 3 – 4 d depending on forage availability. The plots were grazed three times with an average (± standard deviation) regrowth period of 17 (± 6) d. Steers had a similar average daily gain (**ADG**) of 0.87 kg d^-1^ from birth until harvest. Similarly, Bjorklund et al. [19] reported ADG of 0.62 – 0.82 kg d^-1^ for organic grass-fed dairy steers of breeds similar to those used in this study. Steers grazed on WR (0.33 kg d^-1^) and WW (0.32 kg d^-1^) had similar ADG for the grazing period.

### Data collection

#### Weather

The WCROC weather station recorded daily weather. Table 1 reports the monthly mean temperature, total precipitation, and total snowfall for the 130-year long-term mean (1886 – 2016) and for the duration of the current study (September 2015 – June 2016). The temperature during the current study was similar to the long-term mean. Precipitation during the spring months (May and June), and total snowfall were about 45% and 35% less than the long-term mean, respectively.

**Table 1.** Monthly weather for the study (2015 – 2016) and long-term mean (1886 – 2016).

|  | Sep | Oct | Nov | Dec | Jan | Feb | Mar | Apr | May | Jun | Mean |
| --- | --- | --- | --- | --- | --- | --- | --- | --- | --- | --- | --- |
| Temperature, °C<sup>a</sup> |  |  |  |  |  |  |  |  |  |  |  |
| Study | 14 | 6 | 0 | -6 | -11 | -8 | 0 | 3 | 10 | 16 | 2 |
| LTM | 15 | 8 | -1 | -9 | -13 | -10 | -3 | 6 | 14 | 19 | 3 |
|  |  |  |  |  |  |  |  |  |  |  | Sum |
| Precipitation, mm<sup>b</sup> |  |  |  |  |  |  |  |  |  |  |  |
| Study | 34 | 40 | 47<br>5 | 27<br>287 | 7<br>112 | 17<br>155 | 16<br>36 | 47<br>36 | 51 | 48 | 333<br>630 |
| LTM | 59 | 47<br>18 | 25<br>127 | 17<br>178 | 18<br>178 | 18<br>188 | 30<br>198 | 58<br>84 | 76<br>3 | 102 | 447<br>973 |

Daily weather collected from the WCROC weather station (Morris, MN, USA).
LTM = 130-year long-term mean
<sup>a</sup>Temperature expressed as monthly means
<sup>b</sup>Preprecipitation expressed as monthly sums; italics indicate snowfall

#### Forage yield and dry matter

Three random forage samples were collected from each group within the paddock prior to grazing by tossing a 0.23 m^2^ quadrat and clipping the forage inside to a height of 5 cm above the soil (25 April – 10 June 2016). The samples were used to determine DM, herbage mass, forage quality, and FAs.

Forage yield was calculated from sample weights before and after drying for 48 h at 60 °C, and by using the equation: forage yield (kg DM ha^-1^) = kg dry sample ÷ 0.23 m^2^ ÷ 0.0001. The DM was calculated by using the equation: DM (% as-fed) = kg dry sample ÷ kg fresh sample × 100. Group means for forage yield and DM measurements were used to obtain a single measurement for each group within paddock and collection date for WR (n = 48) and WW (n = 48).

#### Forage quality and fatty acids

One sample from each group within paddock and collection date was randomly selected for forage quality analysis for WR (n = 48) and WW (n = 48), and a second sample from each group within paddock and collection date was randomly selected for FA analysis for WR (n = 48) and WW (n = 48). Dried forage samples were ground through a 2 mm screen (Model 4, Wiley Mill^®^, Thomas Scientific, Minneapolis, MN, USA) and were stored in WhirlPak^®^ bags before analysis. Forage quality was determined by near-infrared reflectance spectroscopy using standard equations for forage quality characteristics (Rock River Laboratory, Inc., Watertown, WI, USA). Acid detergent fiber (**ADF**) and neutral detergent fiber (**NDF**) were quantified using Ankom™ procedures (Ankom A2000™, Methods 12 and 13) and total tract NDF digestibility (**TTNDFD**) was quantified using in vitro procedures. Minerals and FAs were determined by inductively coupled plasma optical emission spectrometry and gas chromatographic analysis of FA methyl esters (Eurofins BioDiagnostics, Inc., River Falls, WI), respectively. The FA results are reported as a concentration of total fat, and the remaining results are reported as a percent of forage DM.

### Statistical analysis

The MIXED procedure of SAS/STAT^®^ software [20] was used for all statistical analyses. Fixed factor variables were forage, date, and their interaction. Steer group nested in paddock was a random subject effect and date was repeated using the spatial power covariance structure. Results are reported as least squares means and standard errors. To determine the linear effect of date on FAs, separate models were built for each outcome and date was a continuous covariate.

## Results and discussion

### Forage yield and dry matter

Forage yield for WR was greater (*P* < 0.05) than WW (Table 2), which is consistent with results of previous studies [21–24]. However, the actual yield values are inconsistent with previous studies [21–25]. Variation in yield is mainly due to factors, like year, location, and environment, which differ between studies and affect the yield of small grain forages [21,24,26,27]. Little precipitation and lack of irrigation during the growing period [28], lack of fertilizer [29], and the moderate stocking rate for grazing [30] may have contributed to a lower yield than expected.

**Table 2.** Yield, quality, and mineral composition of winter rye and winter wheat forages.

| Measurement<sup>a</sup> | Winter rye | Winter wheat | SE<sup>b</sup> | P-value |  |  |
| --- | --- | --- | --- | --- | --- | --- |
|  |  |  |  | Forage | Date | F x D |
| Yield, kg DM ha<sup>-1</sup> | 2840. | 2571. | 82. | * | *** | *** |
| Dry matter, % as-fed | 21.0 | 23.2 | 0.43 | * | *** |  |
| Crude protein, % DM | 17.6 | 19.3 | 0.30 | ** | *** | * |
| Fat, % DM | 2.67 | 2.41 | 0.017 | *** | *** |  |
| NDF, % DM | 47.8 | 44.9 | 0.35 | *** | *** | *** |
| ADF, % DM | 30.1 | 28.9 | 0.30 | * | *** | * |
| TTNDFD, % NDF | 56.3 | 56.3 | 0.65 |  | *** |  |
| NEg | 0.446 | 0.440 | 0.0035 |  | *** | * |
| NEm | 0.718 | 0.713 | 0.0038 |  | *** | * |
| TDN, % DM | 65.7 | 65.4 | 0.18 |  | *** | * |
| Starch, % DM | 4.08 | 4.22 | 0.12 |  | *** | *** |
| Calcium, % DM | 0.356 | 0.363 | 0.010 |  | *** | ** |
| Phosphorus, % DM | 0.345 | 0.240 | 0.011 | *** | *** | * |
| Potassium, % DM | 2.84 | 2.69 | 0.050 | * | *** | * |
| Magnesium, % DM | 0.145 | 0.136 | 0.0027 | * | *** | * |

Least square means and standard errors for the grazing period (25 April – 10 June 2016).
NDF = neutral detergent fiber; ADF = acid detergent fiber; TTNDFD = total tract neutral detergent fiber digestibility; NEg = net energy for gain; NEm = net energy for maintenance; TDN = total digestible nutrients; F x D = forage and date interaction
<sup>a</sup>Measurements reported as percent of dry matter; dry matter reported as percent of as-fed; TTNDFD reported as percent of NDF
<sup>b</sup>Standard errors are the same for forages
\* $P < 0.05$ ; \*\* $P < 0.001$ ; \*\*\* $P < 0.0001$

In general, WR had greater yield at the start and WW had greater yield in the latter part of the grazing period (Fig 1). These growth trends are consistent with the results of previous studies [21,24,26,31].

**Fig 1.**
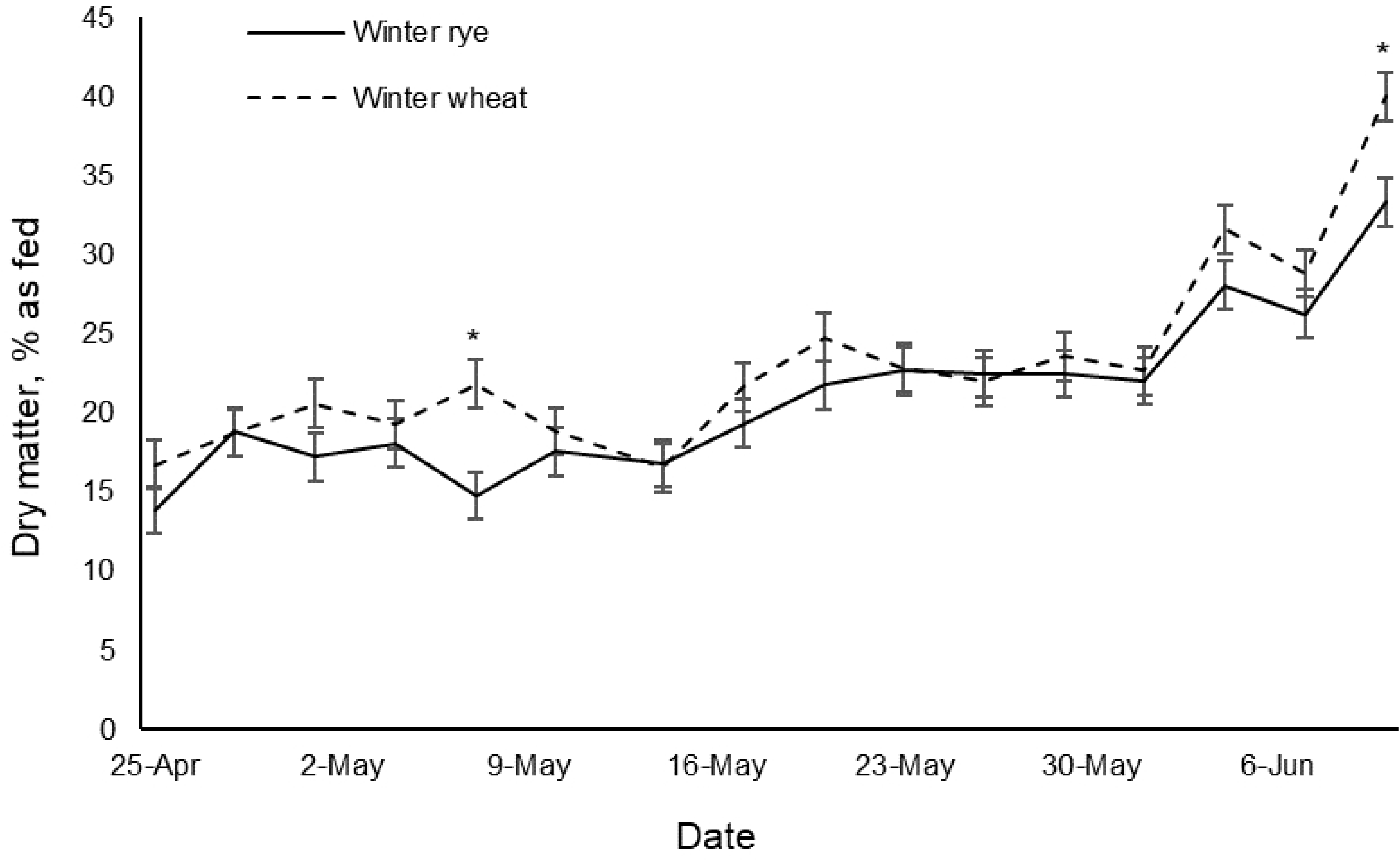
Forage yield of winter rye and winter wheat for the grazing period. *Forages within the same date are different, *P* < 0.05.

The WW had greater (*P* < 0.05) DM than WR (Table 2), which increased during the grazing period for both forages (Fig 2). Previous studies agree with greater DM for WW and an increase in DM during the growing period [23, 28]. The abrupt increase in DM at the end of the grazing period may have been caused by little precipitation in the month of June [28] (Table 1).

**Fig 2.**
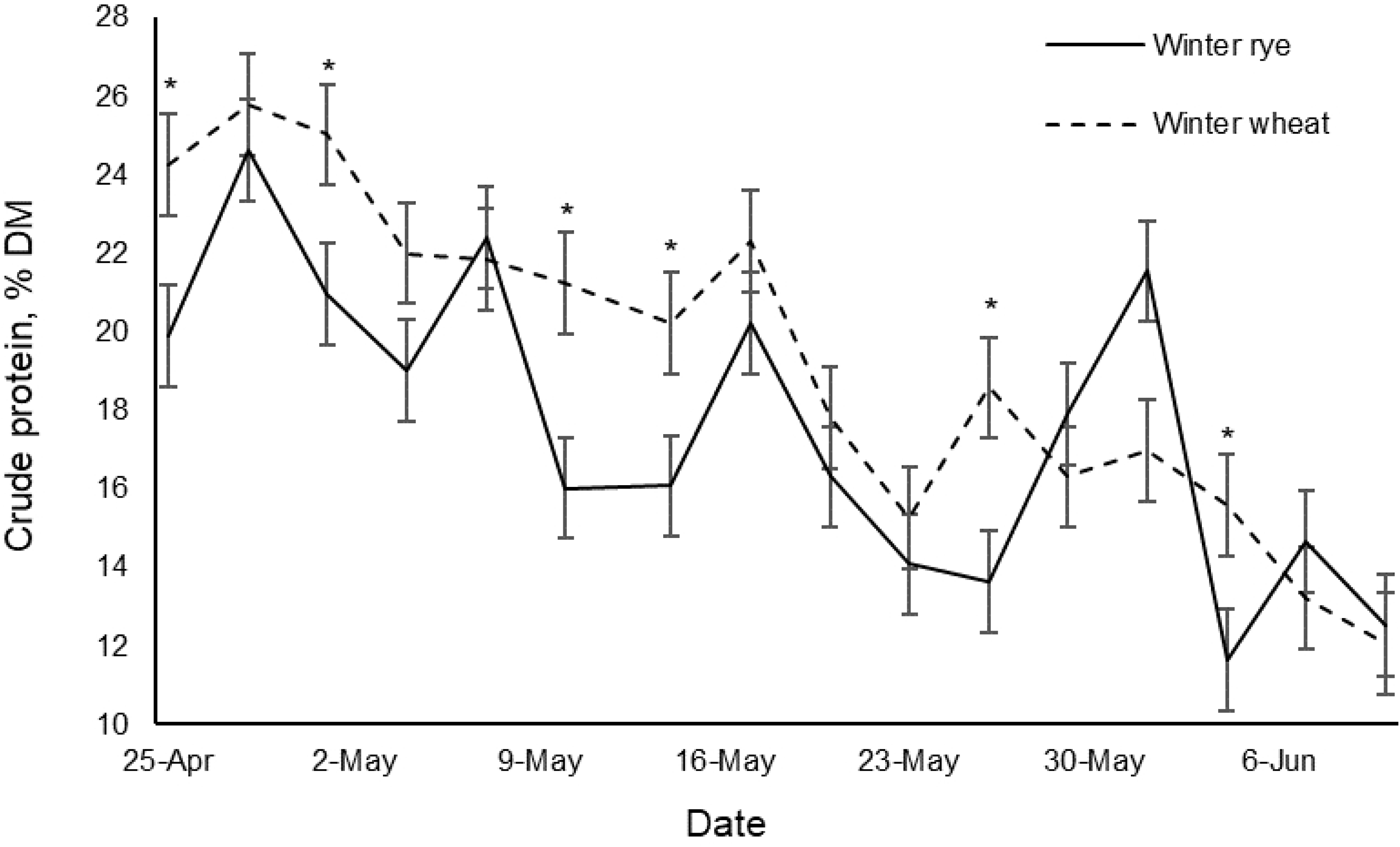
Dry matter of winter rye and winter wheat for the grazing period. *Forages within the same date are different, *P* < 0.05.

### Crude protein and fat

The WW had greater (*P* < 0.001) crude protein (**CP**) than WR (Table 2), which decreased over the grazing period (Fig 3). Previous studies agree with greater CP for WW than WR [21,26,27] and a general decrease in CP over the growing period [15,25–28,32]. The CP values are consistent with previous studies performed in the Midwest region of the USA, which reported CP of 12 – 20% DM for WR [21, 26] and 11 – 34% DM for WW [22,25,26].

**Fig 3.**
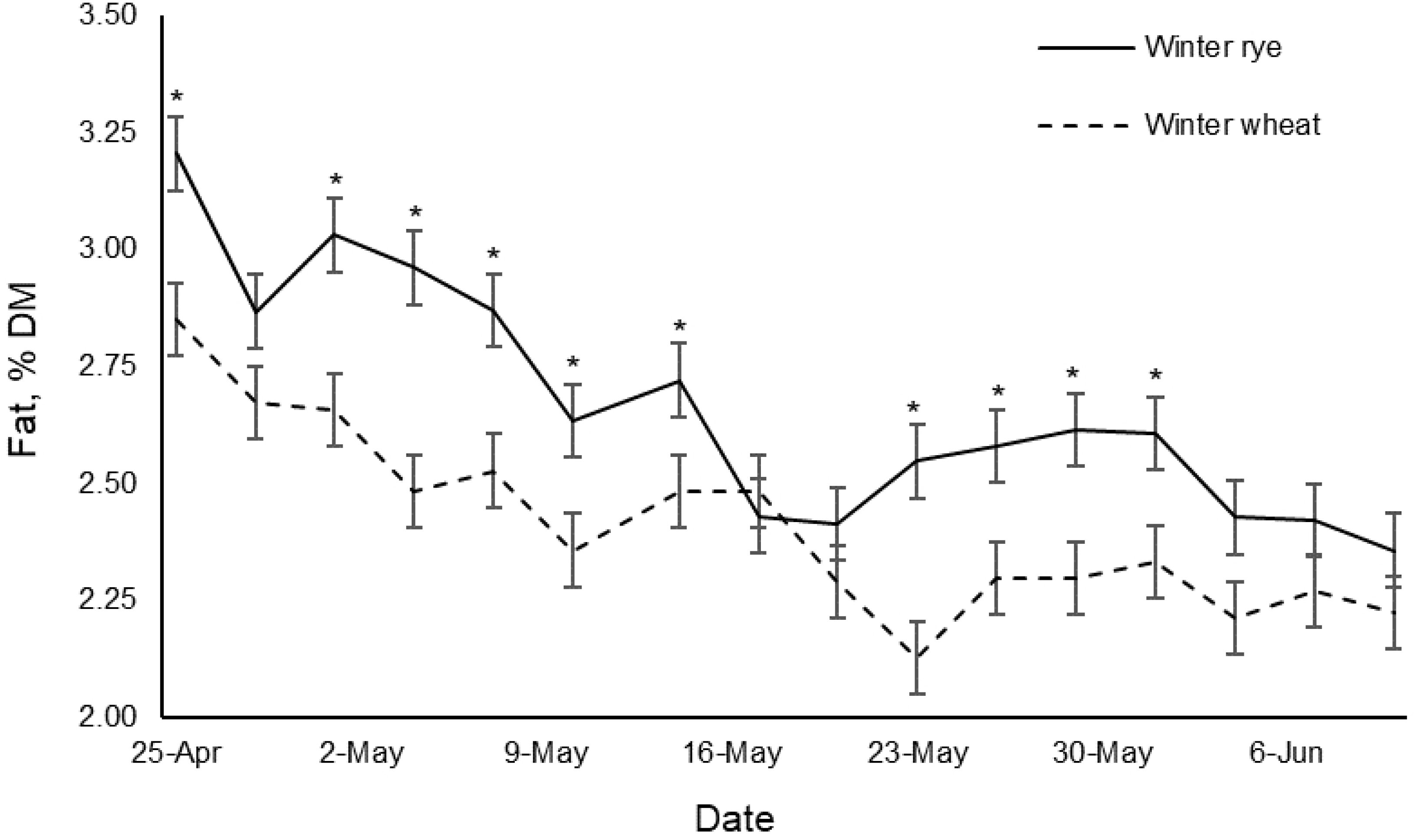
Crude protein of winter rye and winter wheat for the grazing period. *Forages within the same date are different, *P* < 0.05.

The CP levels were well above the minimum requirement for cattle. The estimated minimum CP requirement of 12% DM for growing and finishing beef cattle is based off the estimated net protein requirement for gain of dairy steers consuming forages of the current study with ADG of 0.87 kg d^-1^ using equations from the National Research Council [33]. For dairy cattle, the National Research Council [34] suggests that maximum milk production is observed at 23% DM of CP, which is similar to the values of forages in the first weeks of grazing. The WW forage may be preferred for maximizing milk production. Because CP decreases over the grazing period, early grazing should be initiated to maximize protein intake.

The WR had greater (*P* < 0.0001) fat than WW (Table 2), which decreased over the grazing period for both forages (Fig 4). These results are similar to results by Glasser et al. [9], who reported a decrease in fat from fresh pasture forages as cutting date increased.

**Fig 4.**
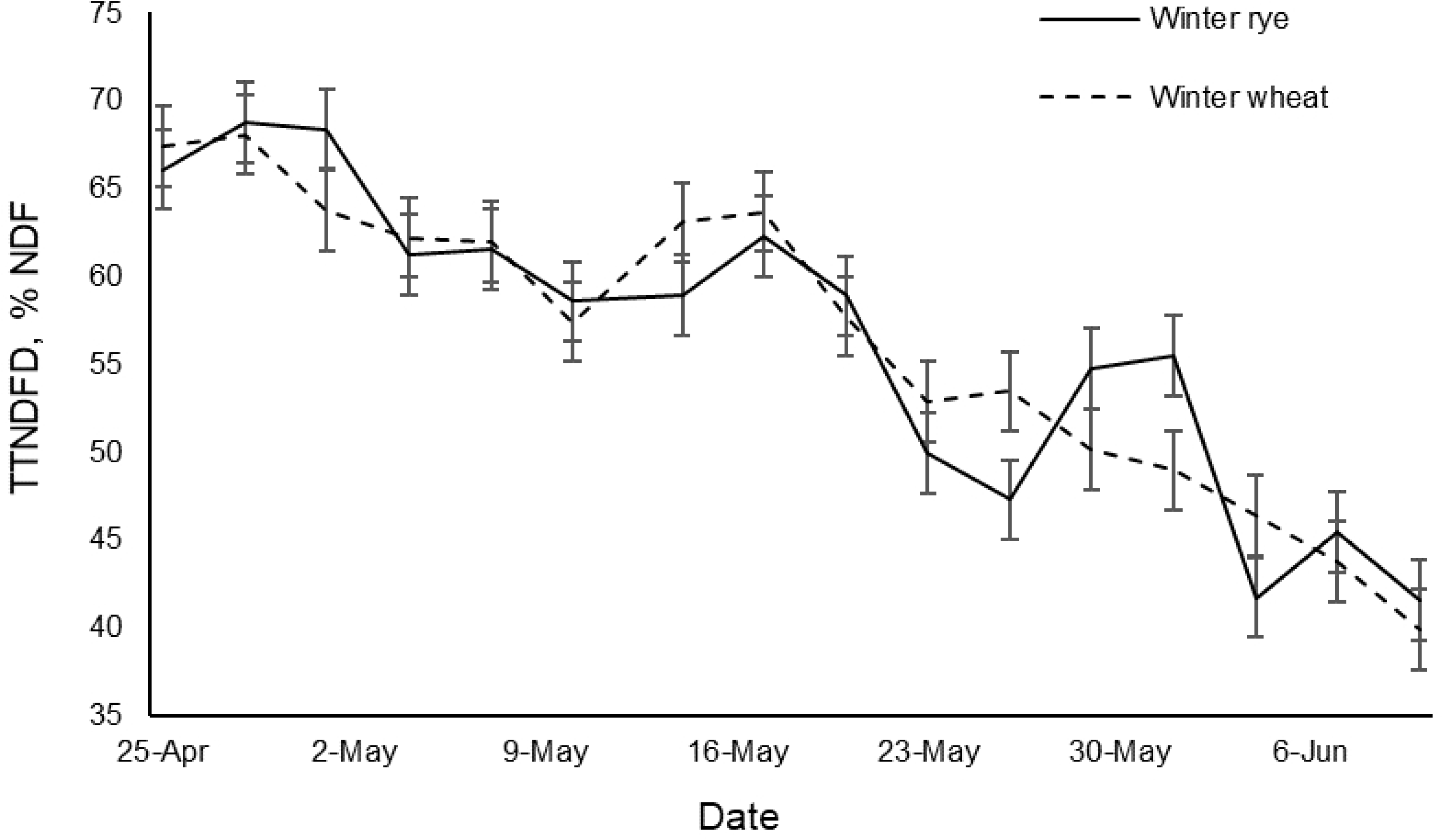
Fat of winter rye and winter wheat for the grazing period. *Forages within the same date are different, *P* < 0.05.

### Fiber and digestibility

#### Neutral detergent fiber and acid detergent fiber

The WR had greater (*P* < 0.05) NDF and ADF compared to WW (Table 2), and forages increased in NDF and ADF over the grazing period. Previous studies reported greater fiber for WR compared to WW [21, 27] and an increase in NDF and ADF with plant maturity [15,25,27,28]. Similar studies performed in the Midwest region of the USA [21, 25] reported NDF of 53 – 67% DM for WR and 38 – 61% DM for WW, and ADF of 28 – 42% DM for WR and 24 – 31% DM for WW. Low precipitation during the current study may have slowed maturation and decreased fiber production [28]. The forages had similar NDF values at the end of the grazing period, which is similar to findings by Lauriault and Kirksey [31]. Because research is sparse, the National Research Council does not have specific recommendations for NDF or ADF of diets for grazing cattle [34].

#### Total tract neutral detergent fiber digestibility

The forages had similar TTNDFD and met the minimum recommended value of 50% NDF [35], except during the last two weeks of the grazing period (Fig 5). The decrease in digestibility during the growing period agrees with the results of previous studies [15,26,28]. Grazing mature WR and WW forages may not meet TTNDFD recommendations, but they might still be adequate for grazing.

**Fig 5.**
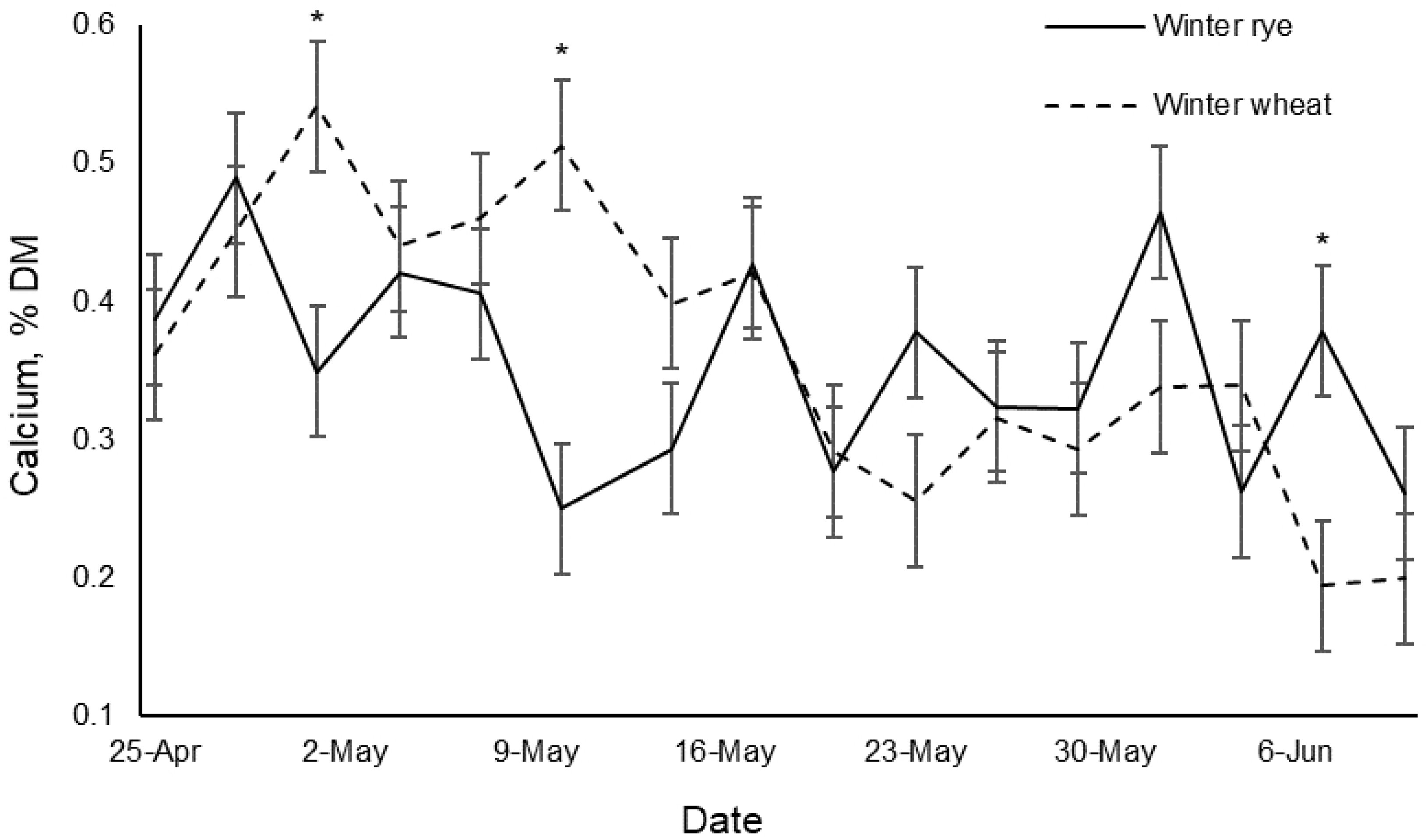
TTNDFD of winter rye and winter wheat for the grazing period. TTNDFD = total tract neutral detergent fiber digestibility

### Minerals

#### Calcium

The forages had similar Calcium (**Ca**) (Table 2), which decreased over the grazing period (Fig 6). The estimated Ca requirement of 0.40 – 0.5% DM for growing and finishing beef cattle is based off the estimated net protein requirement for gain of dairy steers consuming forages of the current study with ADG of 0.87 kg d^-1^ using equations from the National Research Council [33, 36]. The forages met the Ca requirements for beef cattle in the first half, but were deficient in the latter half of the grazing period. For dairy cattle, the National Research Council [34] recommends absorbed Ca of 1.22 – 1.45 g kg^-1^ of milk produced. Since the typical organic dairy cow in the USA produces approximately 20 kg d^-1^ of milk [37–39] and Ca is approximately 30% available for absorption in forages [34], grazing dairy cattle need to consume an estimated 83.4 – 99.1 g d^-1^ of Ca from pasture forages. Vazquez et al. [40] estimated that a grazing dairy cow consumes 8.7 – 14.6 kg DM d^-1^ of forage. Therefore, the estimated Ca requirement in pasture forages for grazing dairy cattle is 0.96 – 1.15% DM. The forages of the current study were deficient in Ca for dairy cattle, so supplemental Ca for grazing lactating dairy cattle is necessary.

**Fig 6.**
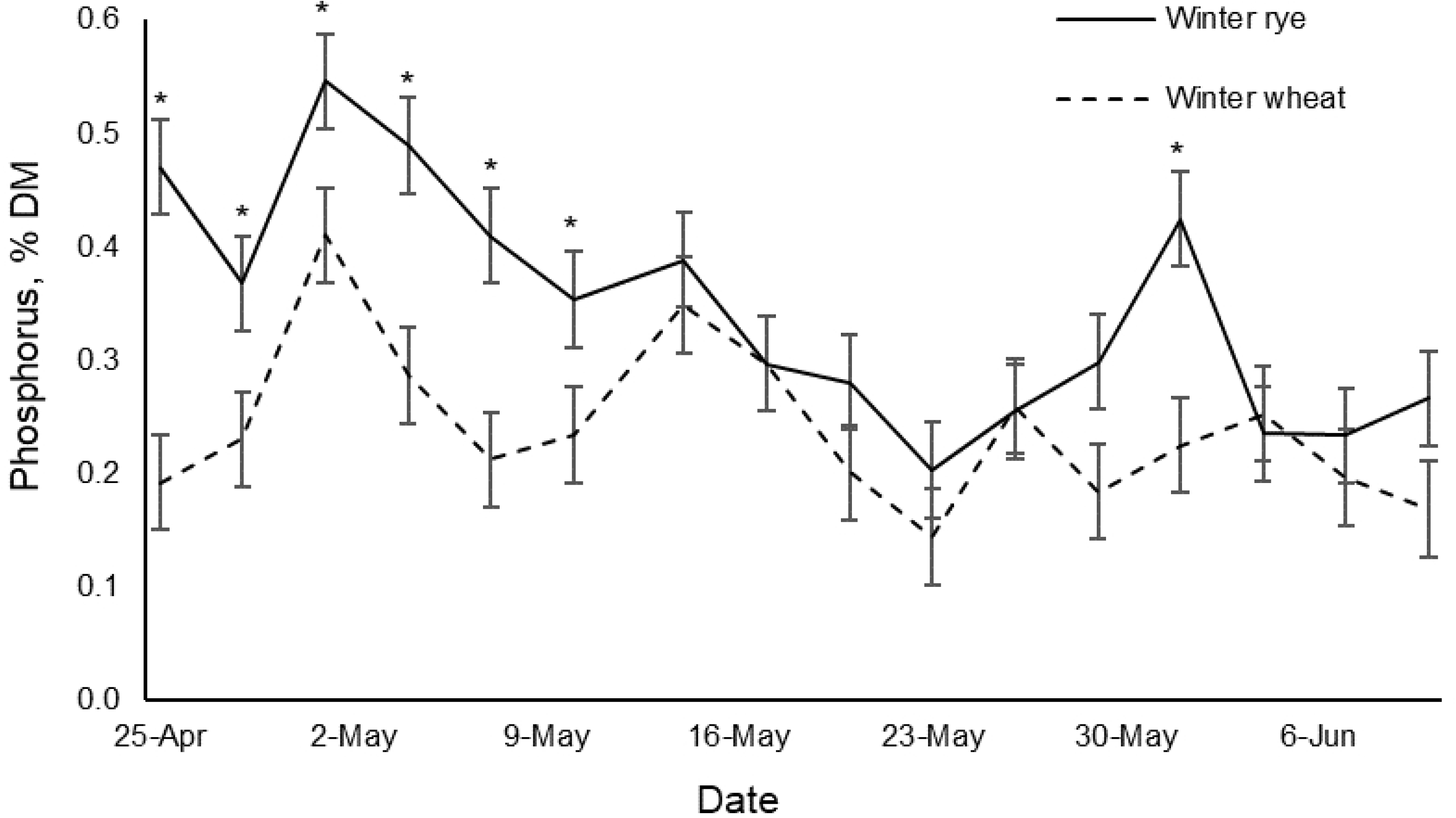
Calcium of winter rye and winter wheat for the grazing season. *Forages within the same date are different, *P* < 0.05.

#### Phosphorus

The WR had greater (*P* < 0.0001) phosphorus (**P**) than WW (Table 2), which decreased over the grazing period (Fig 7). Specifically, WR had greater P in the first few weeks of grazing compared to WW. The estimated P requirement of 0.16 – 0.21% DM for growing and finishing beef cattle is based off the estimated net protein requirement for gain of dairy steers consuming forages of the current study with ADG of 0.87 kg d^-1^ using equations from the National Research Council [33, 36]. The forages were above the minimum P requirements for beef cattle and did not reach the maximum tolerable level of 0.7% DM during the grazing period [33]. However, the forages were lower than the National Research Council [34] recommendation of 0.32% DM for P estimated for lactating grazing dairy cattle [37–40], especially in the latter half of grazing. Therefore, supplemental P for lactating dairy cattle is necessary for grazing WW and for WR after the first few weeks of grazing.

**Fig 7.**
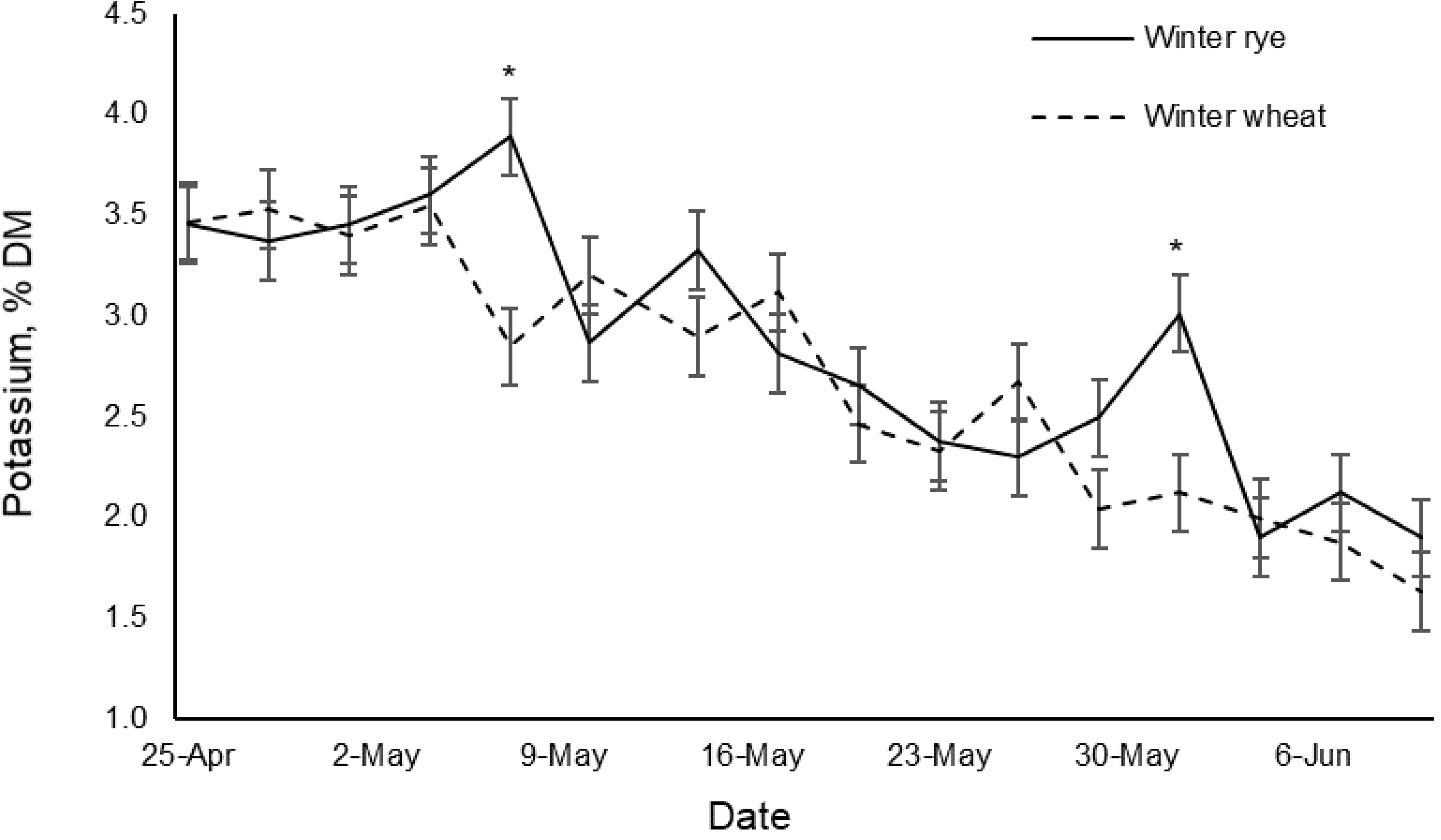
Phosphorus of winter rye and winter wheat for the grazing period. *Forages within the same date are different, *P* < 0.05.

#### Potassium

The WR had greater (*P* < 0.05) potassium (**K**) than WW (Table 2), which decreased during the grazing period (Fig 8). In general, the forages were well above the National Research Council [33, 34] recommendations of 0.42% DM for lactating grazing dairy cattle and 0.60% DM for growing beef cattle, and was above the maximum tolerable level of 2 – 3% DM during the beginning of the grazing period. The forages exceeded the maximum tolerable level at the start of grazing, which is a concern for lactating dairy cattle. High levels of K may reduce dry matter intake, milk yield, and inhibit Ca and magnesium (**Mg**) absorption [34, 36]. Therefore, supplemental K is likely unnecessary for cattle grazing WR and WW forages.

**Fig 8.**
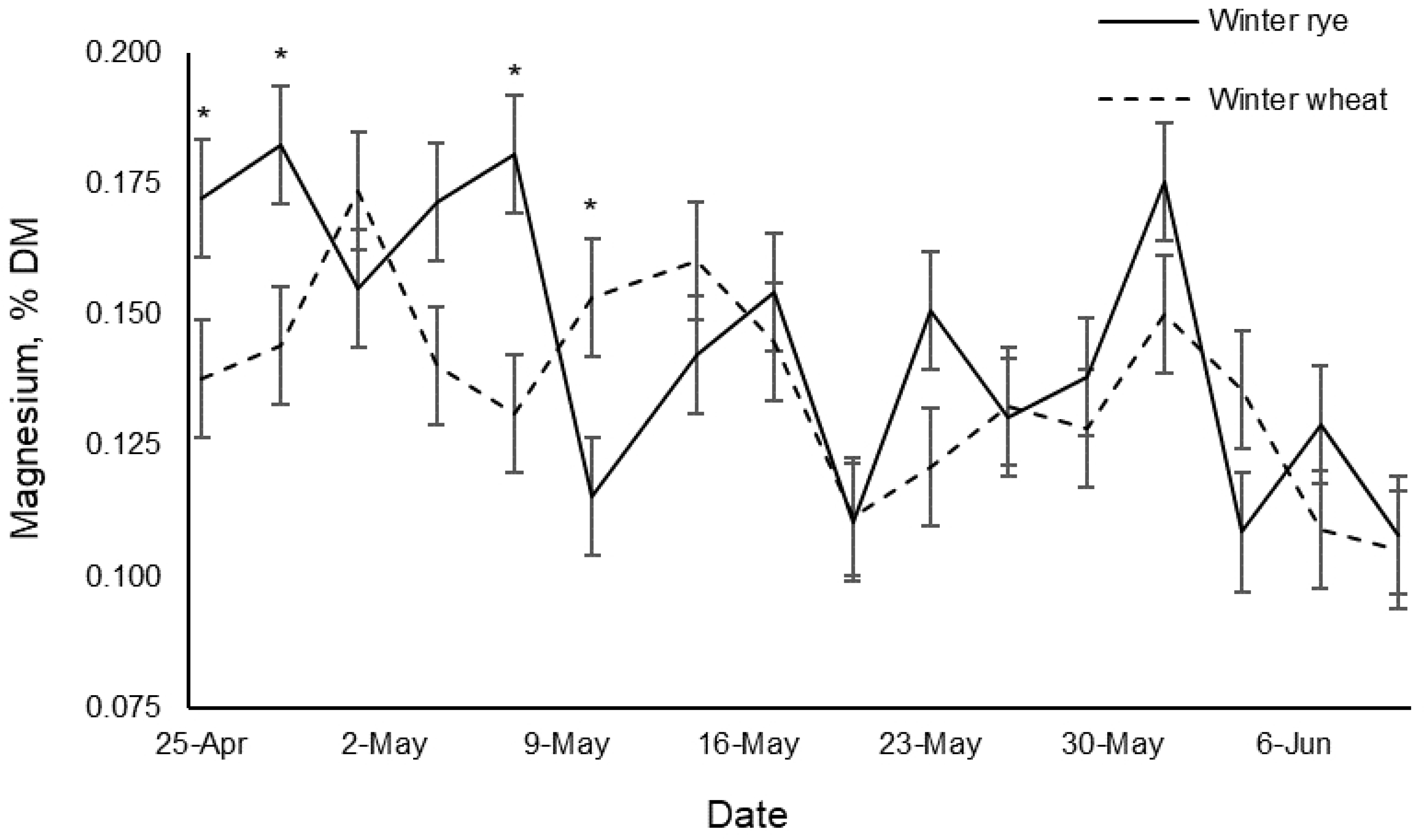
Potassium of winter rye and winter wheat for the grazing period. *Forages within the same date are different, *P* < 0.05.

#### Magnesium

The WR had greater (*P* < 0.05) Mg than WW (Table 2), which decreased over the grazing period (Fig 9). Mg deficiency (hypomagnesemia) is a concern for pastures with rapidly growing cereal crops in the early spring [36]. Dove et al. [36] suggested that hypomagnesemia may also be induced by a K:(Mg + Ca) ratio greater than 2.2. Forages met the Mg requirement for beef cattle of 0.10 – 0.20% DM [33, 36] for the grazing period; however, the forages were lower than the recommended Mg levels for lactating dairy cattle of 0.35 – 0.40% DM [34, 41]. Therefore, supplemental Mg for lactating dairy cattle is necessary for grazing WR and WW forages.

**Fig 9.**
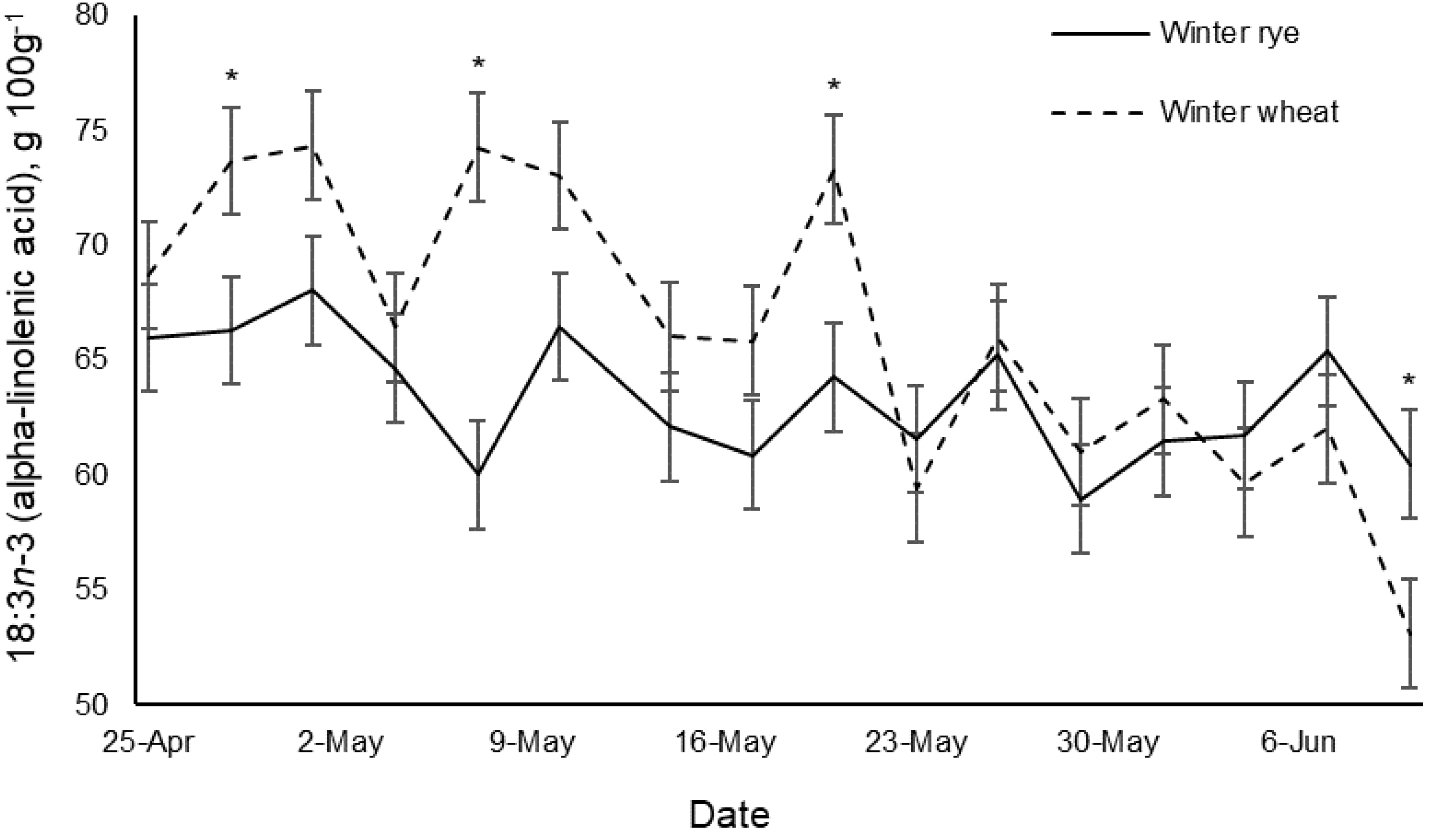
Magnesium of winter rye and winter wheat for the grazing period. *Forages within the same date are different, *P* < 0.05.

#### Fatty acids

The FAs for WR and WW forages differed (*P* < 0.05) in myristic acid (**14:0**), linoleic acid (**18:2*n*-6**), 18:3*n*-3, arachidic acid (**20:0**), eicosenoic acid (**20:1**), behenic acid (**22:0**), and lignoceric acid (**24:0**) (Table 3). The most abundant was 18:3*n*-3, followed by palmitic acid (**16:0**) and 18:2*n*-6.

**Table 3.**
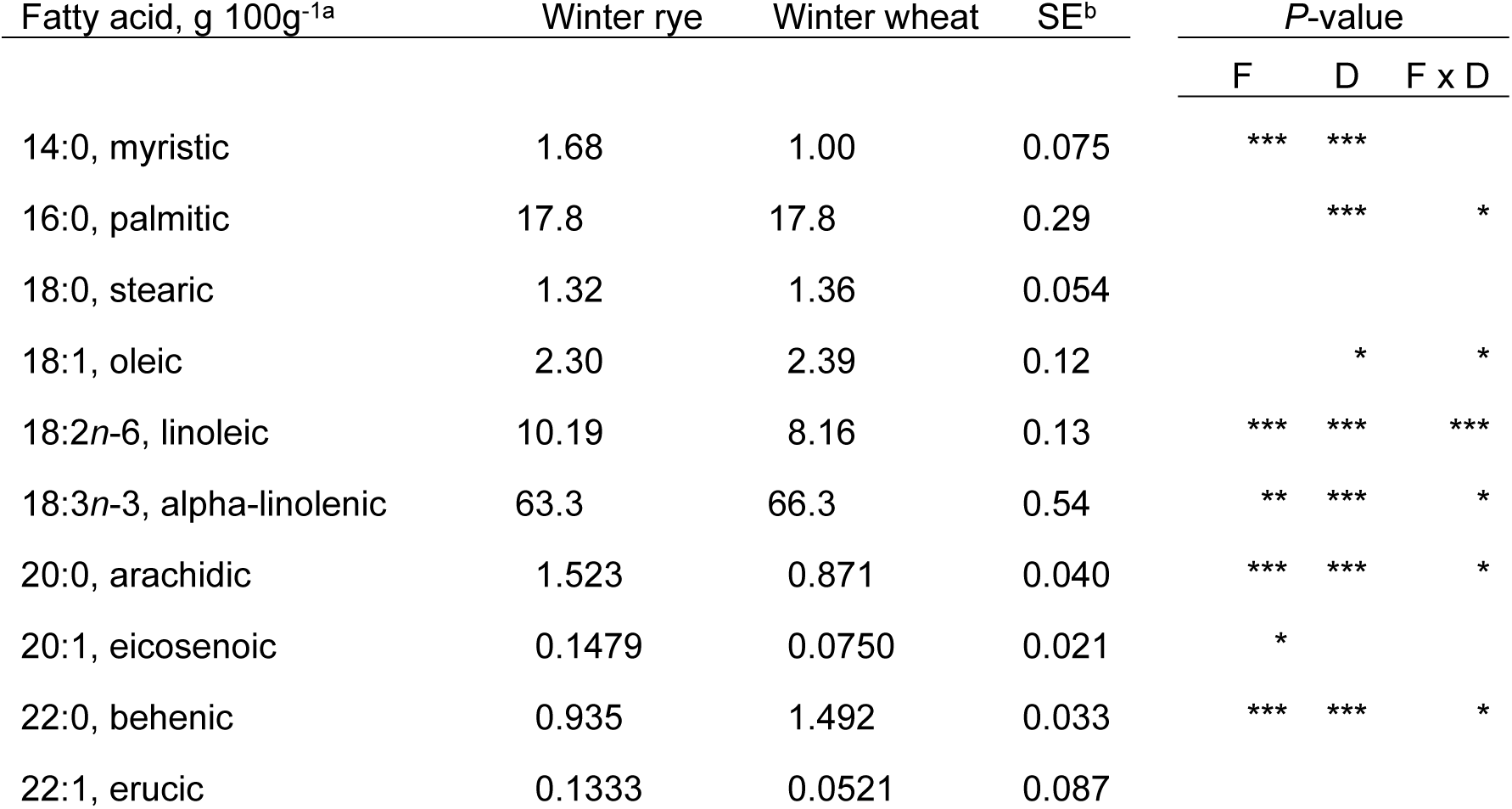
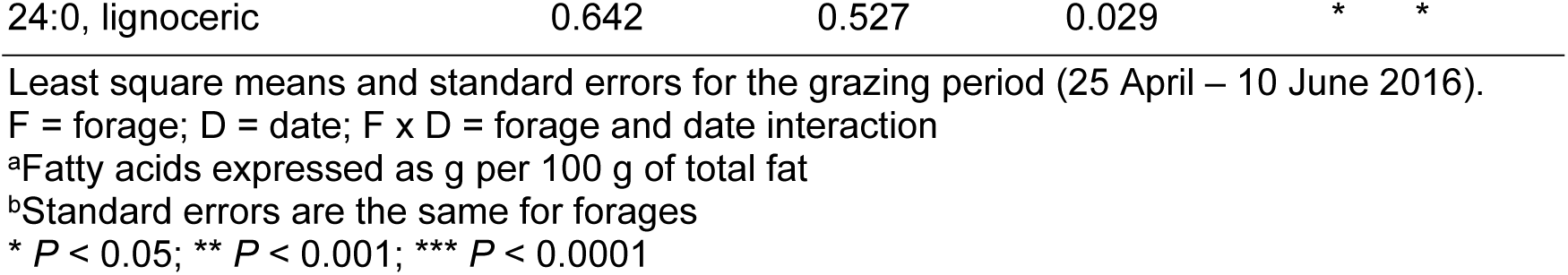
Fatty acids of winter rye and winter wheat forages.

| Fatty acid, g 100g<sup>-1a</sup> | Winter rye | Winter wheat | SE<sup>b</sup> | P-value |  |  |
| --- | --- | --- | --- | --- | --- | --- |
|  |  |  |  | F | D | F x D |
| 14:0, myristic | 1.68 | 1.00 | 0.075 | *** | *** |  |
| 16:0, palmitic | 17.8 | 17.8 | 0.29 |  | *** | * |
| 18:0, stearic | 1.32 | 1.36 | 0.054 |  |  |  |
| 18:1, oleic | 2.30 | 2.39 | 0.12 |  | * | * |
| 18:2*n*-6, linoleic | 10.19 | 8.16 | 0.13 | *** | *** | *** |
| 18:3*n*-3, alpha-linolenic | 63.3 | 66.3 | 0.54 | ** | *** | * |
| 20:0, arachidic | 1.523 | 0.871 | 0.040 | *** | *** | * |
| 20:1, eicosenoic | 0.1479 | 0.0750 | 0.021 | * |  |  |
| 22:0, behenic | 0.935 | 1.492 | 0.033 | *** | *** | * |
| 22:1, erucic | 0.1333 | 0.0521 | 0.087 |  |  |  |

|  |  |  |  |  |  |
| --- | --- | --- | --- | --- | --- |
| 24:0, lignoceric | 0.642 | 0.527 | 0.029 | * | * |

Least square means and standard errors for the grazing period (25 April – 10 June 2016).
F = forage; D = date; F x D = forage and date interaction
<sup>a</sup>Fatty acids expressed as g per 100 g of total fat
<sup>b</sup>Standard errors are the same for forages
\* $P < 0.05$ ; \*\* $P < 0.001$ ; \*\*\* $P < 0.0001$

Beef lipid FAs of this study are presented in Phillips et al. [18]. Briefly, beef lipids from steers grazed on WR and WW differed (*P* < 0.05) in butyric acid (4:0), tetradecenoic acid (14:1*trans*), myristoleic acid (14:1), hexadecenoic acid (16:1*trans*), margaroleic acid (17:1), octadecadienoic acid (18:2*trans*), gamma-linolenic acid (18:3*n*-6), eicosatrienoic acid (20:3*n*-3), arachidonic acid (20:4*n*-6), heneicosanoic acid (21:0), and docosadienoic acid (22:2*n*-6). The sum of saturated, *cis*-monounsaturated, *cis*-polyunsaturated, *trans*, and *n*-6 fats were similar between steers. The steers grazed on WR (0.535 ± 0.018 g 100g^-1^) had numerically less (*P* = 0.31) *n*-3 than steers grazed on WW (0.562 ± 0.018 g 100g^-1^). The differences in FAs of WR and WW forages may have contributed to differences in beef lipid FAs of steers [7]. However, the duration of grazing forages (7 weeks) may not have been long enough to investigate the effects of forages and their contributions to individual FA levels of beef lipids.

Table 4 presents the estimated linear effect of date for FAs during the grazing period for WR and WW forages. For Table 4, significance for WR or WW terms indicates a linear effect of date, whereas significance for the interaction term indicates a difference in the effect of date between forages. Thus, the 14:0, erucic acid (**22:1**), and 24:0 increased (*P* < 0.05) during the grazing period, and the effect of date was similar for forages. The effect of date differed (*P* < 0.05) between forages for 16:0, oleic acid (**18:1**), 18:2*n*-6, 18:3*n*-3, and 20:0. The 18:1 increased (*P* < 0.0001) in WW, but did not change during the grazing period for WR. The 20:0 decreased 1.41 times more (*P* < 0.05) per d for WR compared to WW. There was no effect of date for stearic acid (**18:0**), 20:1, and 22:0 during the grazing period.

**Table 4.**
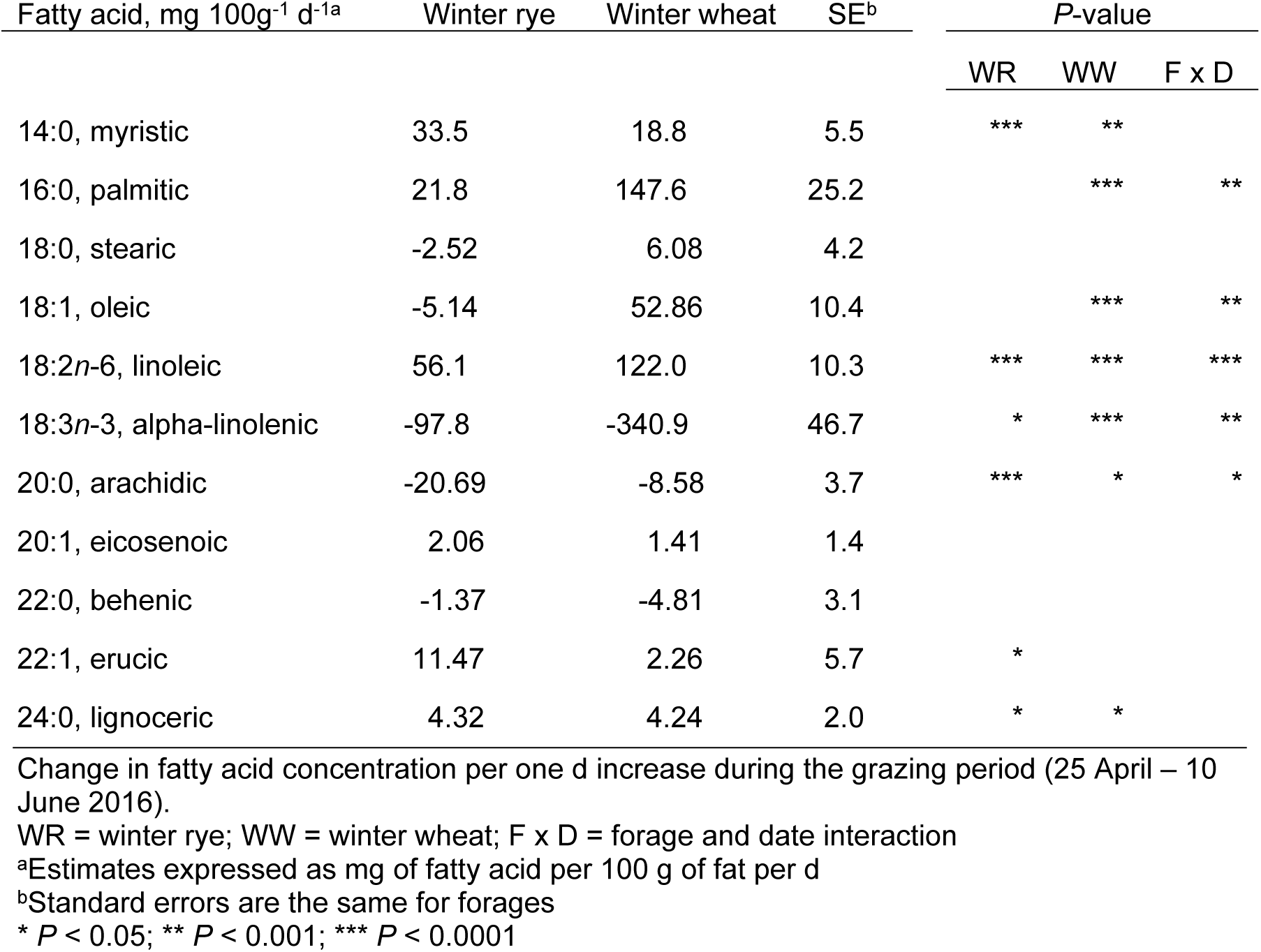
Effect of date on fatty acids of winter rye and winter wheat forages.

#### Alpha-linolenic acid, 18:3*n*-3

The WW was 4.7% greater (*P* < 0.001) in 18:3*n*-3 concentration compared to WR. The 18:3*n*-3 in WW decreased 2.49 times more (*P* < 0.001) per d compared to WR (Table 4). Clapham et al. [42] reported 18:3*n*-3 of 65 – 69 g 100g^-1^ for triticale (a hybrid of rye [*Secale*] and wheat [*Triticum*]), which is similar to values for WW of the current study. The study [42] also reported a general decrease in 18:3*n*-3 with plant maturity, which is similar to forages of the current study. In a meta-analysis, Glasser et al. [9] reported a decrease in 18:3*n*-3 concentration as pasture forages matured. Fig 10 depicts the decrease in 18:3*n*-3 for forages during the grazing period.

**Fig 10.**
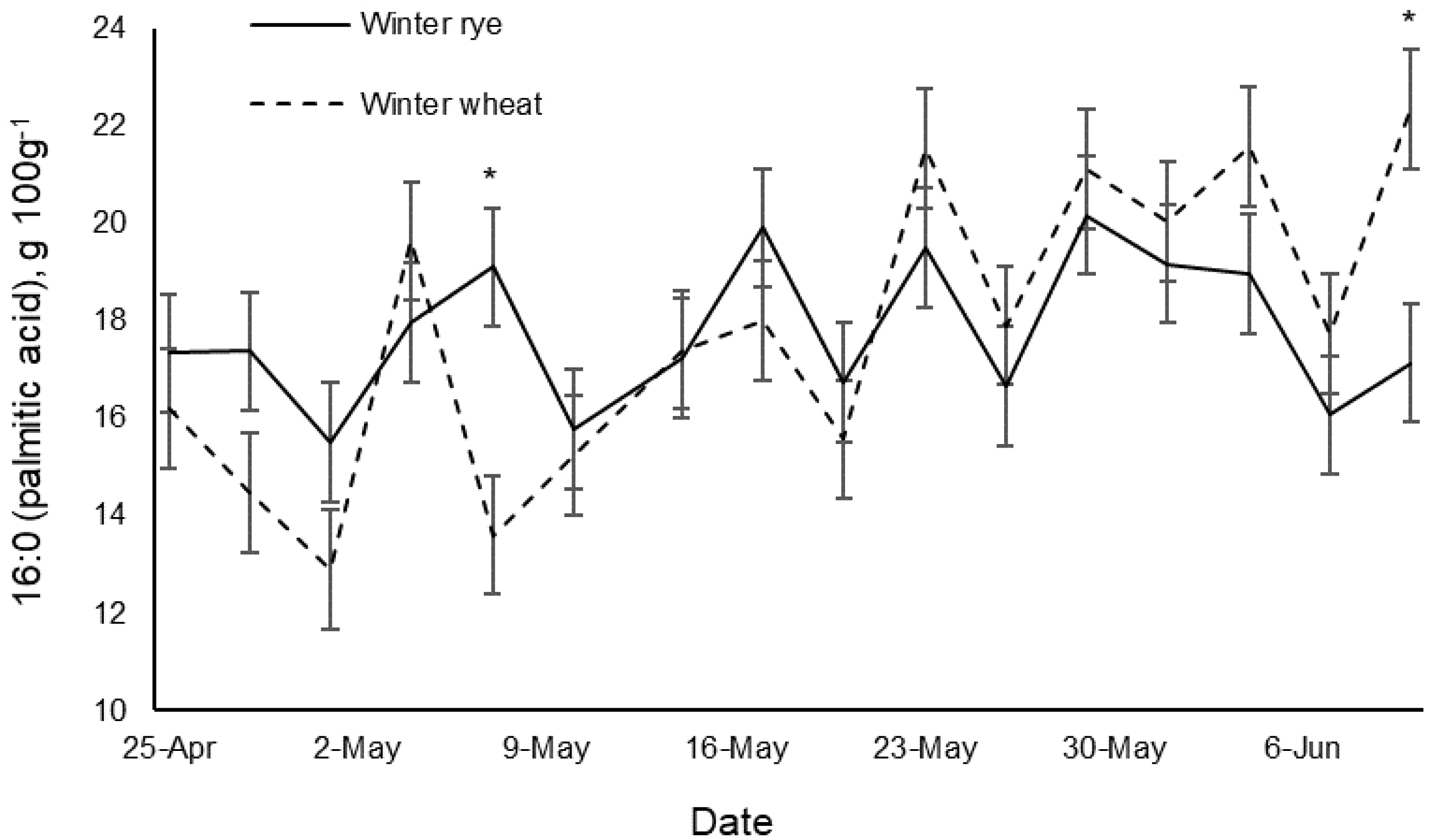
Alpha-linolenic acid (18:3*n*-3) of winter rye and winter wheat for the grazing period. *Forages within the same date are different, *P* < 0.05.

#### Palmitic acid, 16:0

The forages had similar 16:0 concentration. The 16:0 increased (*P* < 0.0001) in WW; however, there was no effect of date for WR (Table 4). That is, the 16:0 increased in WW and did not radically change in WR during the grazing period. Clapham et al. [42] reported lower 16:0 of 13 – 15 g 100g^-1^ for triticale. The lack of additional nitrogen fertilizer used in the current study may have increased the 16:0 in forages [9]. In a meta-analysis, Glasser et al. [9] reported an increase in 16:0 concentrations as pasture forages matured. Temporal changes in 16:0 are illustrated in Fig 11.

**Fig 11.**
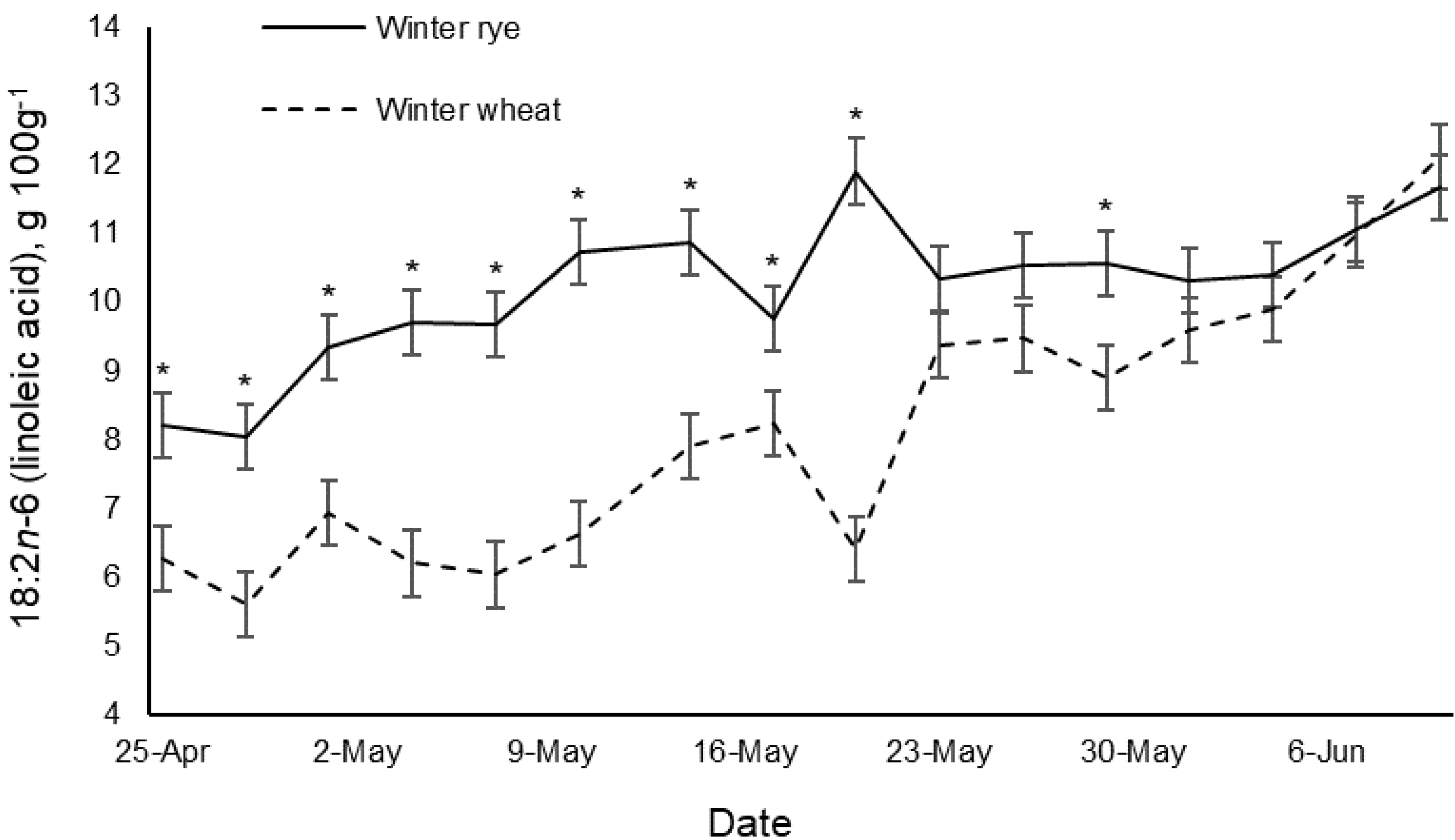
Palmitic acid (16:0) of winter rye and winter wheat for the grazing period. *Forages within the same date are different, *P* < 0.05.

#### Linoleic acid, 18:2*n*-6

The WR had 24.9% greater (*P* < 0.0001) 18:2*n*-6 concentration than WW. The 18:2*n*-6 increased 1.17 times more (*P* < 0.0001) per d for WW compared to WR (Table 4). Clapham et al. [42] reported greater 18:2*n*-6 values of 12 – 13 g 100g^-1^ and a similar increase in 18:2*n*-6 as triticale matured. Fig 12 illustrates the increase in 18:2*n*-6 for forages during the grazing period. The forages differed in the first half of grazing, but were similar in the latter half of grazing.

**Fig 12.**
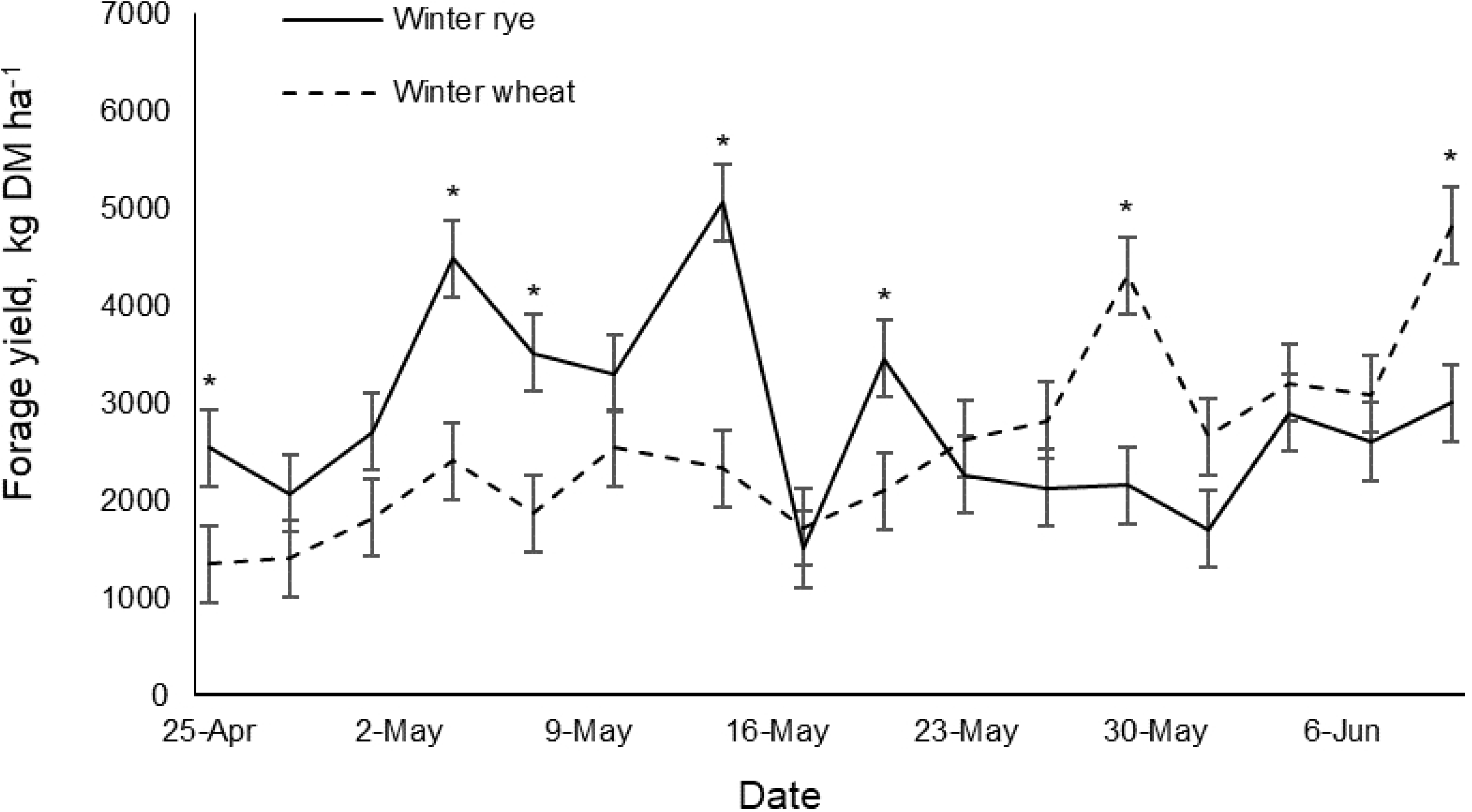
Linoleic acid (18:2*n*-6) of winter rye and winter wheat for the grazing period. *Forages within the same date are different, *P* < 0.05.

### Comparison to perennial pasture forages

The grazing season for perennial pastures in the upper Midwest of the USA is late-May – October. The small grain forages of the current study extended the grazing season by approximately one month earlier. A study by Ruh [43] conducted at the WCROC assessed the forage quality of cool season perennial forages during the grazing season (June – October 2017). Ruh [43] reported 343 – 612 kg DM ha^-1^ less in average yield and 3.7 – 5.4% DM greater CP for perennial forages compared to the small grain forages of the current study. The study [43] also reported greater NDF (49.6% DM), ADF (32.2% DM), and lower TTNDFD (54.6% NDF) for perennial pastures, and greater Ca (0.67% DM), K (3.10% DM), and Mg (0.23% DM) compared to the small grain forages of the current study. Perennial forages had similar P (0.33% DM) compared to WR, and greater P compared to WW of the current study. Therefore, the vegetative stages of WR and WW pasture forages may provide an abundance of biomass and similar nutritional quality as perennial pastures for grazing, while extending the grazing season.

A meta-analysis of FAs in grasses performed by Glasser et al. [9] reported means for 16:0 (16.9 g 100g^-1^), 18:2*n*-6 (15.8 g 100g^-1^), and 18:3*n*-3 (52.6 g 100g^-1^) in alfalfa, red clover, white clover, and multi-species mixed grasses. Glasser et al. [9] reported similar 16:0 (16.9 g 100g^-1^), greater 18:2*n*-6 (15.8 g 100g^-1^), and lower 18:3*n*-3 (52.6 g 100g^-1^) than forages of the current study. The greater 18:3*n*-3 found in the WR and WW of the current study is preferred since dietary 18:3*n*-3 increases *n*-3 FAs in the meat and milk of cattle. Therefore, grazing cattle on WR and WW forages may even enhance the FA profiles of beef and dairy products, potentially benefitting producers and consumers.

## Conclusions

Based on results from this study and the previous supplemental study by Phillips et al. [18], winter rye and winter wheat forages are viable options for cattle grazing in the early spring and summer in the Midwest of the USA. Results suggested that winter rye might offer more herbage mass in the early spring at the expense of lower crude protein and 18:3*n*-3 concentration compared to winter wheat. The greater 18:3*n*-3 concentration for winter wheat may contribute to healthier beef and milk fatty acids. In general, crude protein, digestibility, minerals, and alpha-linolenic acid decreased, and fiber and linoleic acid increased during the grazing period. Therefore, results of this study suggest that producers should initiate early grazing in the spring to maximize digestibility, protein, and alpha-linolenic acid intake while the small grain forages are immature. To mitigate mineral imbalance, free-choice minerals formulated for pasture grazing must be offered to meet mineral demands. With fatty acids of beef and dairy products becoming a critical part of consumer choice and health, this study also showed that alpha-linolenic acid is abundant in winter rye and winter wheat forages, which could contribute to healthier milk and beef products.

## Acknowledgements

The authors express gratitude to Darin Huot and staff at WCROC for their assistance in data collection and care of animals. This project was supported by the National Institute of Food and Agriculture, United States Department of Agriculture, under award number 2014-51300-22541.

